# 3D super-resolution fluorescence microscopy maps the variable molecular architecture of the Nuclear Pore Complex

**DOI:** 10.1101/2020.11.27.386599

**Authors:** Vilma Jimenez Sabinina, M. Julius Hossain, Jean-Karim Hériché, Philipp Hoess, Bianca Nijmeijer, Shyamal Mosalaganti, Moritz Kueblbeck, Andrea Callegari, Anna Szymborska, Martin Beck, Jonas Ries, Jan Ellenberg

## Abstract

Nuclear pore complexes (NPCs) are large macromolecular machines that mediate the traffic between the nucleus and the cytoplasm. In vertebrates, each NPC consists of ~1000 proteins, termed nucleoporins, and has a mass of over 100 MDa. While a pseudo-atomic static model of the central scaffold of the NPC has recently been assembled by integrating data from isolated proteins and complexes, many structural components still remain elusive due to the enormous size and flexibility of the NPC. Here, we explored the power of 3D super-resolution microscopy combined with computational classification and averaging to explore the 3D structure of the NPC in single human cells. We show that this approach can build the first integrated 3D structural map containing both central as well as peripheral NPC subunits with molecular specificity and nanoscale resolution. Our unbiased classification of over ten thousand individual NPCs indicates that the nuclear ring and the nuclear basket can adopt different conformations. Our approach opens up the exciting possibility to relate different structural states of the NPC to function in situ.

## Introduction

The Nuclear Pore Complex (NPC) is the gateway for the tightly regulated exchange of macromolecules between the nucleus and the cytoplasm and is one of the largest protein complexes in eukaryotic cells with an estimated molecular mass of over 100 MDa. Each NPC is formed by multiple copies of around 30 different nucleoporins (Nups) (Cronshaw et al., 2002), modularly arranged into about seven distinct subcomplexes (Schwartz, 2005) with an eightfold rotational symmetry. The central scaffold of the NPC is composed of three stacked, ring-shaped structures called the inner, cytoplasmic and nucleoplasmic rings. In addition, the cytoplasmic filaments and the nuclear basket, which are localized asymmetrically, complete the NPC’s peripheral architecture. Although an integrated pseudo-atomic model of the central scaffold of the mature vertebrate NPC has been proposed by computationally integrating data of isolated Nups and NPCs from a static perspective (reviewed in Hampoelz et al., 2019; Lin and Hoelz, 2019) such atomic resolution methods face challenges when assessing asymmetrically positioned central Nups, and peripheral components of the NPC assumed to be present in variable conformational states.

Super-resolution microscopy using single molecule localization (SMLM) has previously allowed the direct visualization of different components of the NPC scaffold (Göttfert et al., 2013; Loschberger et al., 2012; Ori et al., 2013; Thevathasan et al., 2019) and its combination with 2D particle averaging retrieved precise measurements of the radial localization of scaffold Nups (Heydarian et al., 2018; Szymborska et al., 2013). In this study, we implemented high precision two-color 3D SMLM with data analysis by particle averaging and unsupervised classification to map the major NPC substructures including its cytoplasmic and nuclear filaments, in human cells. To this end, we generated a set of genome-edited cell lines with fluorescently tagged nucleoporins for major building blocks of the pore, the Y-shaped complex that forms the nuclear and cytoplasmic rings (SEH1, NUP107, NUP133), the nuclear basket (TPR) and the cytoplasmic filaments (RANBP2/NUP358).

Our quantitative integrated imaging and computational methodology allowed us to map the 3D distribution of all individual Nups, clearly revealing the size and positions of differently sized double and single rings along the transport axis and an unexpectedly large nuclear volume explored by TPR. Integrating the data of all five Nups by particle averaging of the common ELYS reference Nup then allowed us to create the first multimolecular 3D structure model of the NPC based on light microscopy that provides novel information for three structural components of the NPC. Exploring the distribution of ring geometry among individual super-resolved NPCs furthermore suggested the presence of different conformational states. Indeed, unsupervised clustering of the 3D point cloud localizations of all single pore particles using topological data analysis revealed different classes of NPCs for individual Nups. Comparison of their structural differences by class specific particle averaging revealed potential NPC assembly intermediates for the nuclear ring, and an unexpectedly high degree of variability for the nuclear basket.

## Results

In this study, we systematically imaged the 3D architecture of the major substructures within the NPC with 3D SMLM (Huang et al., 2008; Li et al., 2018) in single human cells. To this end, we tagged the N-terminal domain of several scaffold Nups (SEH1, NUP107 and NUP133), the cytoplasmic filament Nup (NUP358/RANBP2) and the C-terminal domain of the nuclear basket Nup (TPR) (**Fig. 1A**), with a small self-labeling SNAP-tag (Keppler et al., 2003) at all endogenous alleles in U2OS cells by homozygous genome editing (Koch et al., 2018) to allow high efficiency fluorophore labeling with comparable stoichiometry between proteins (Thevathasan et al., 2019). In addition, we used antibody labeling to orthogonally detect the Nup ELYS in a second color, as a constant reference to determine the relative position of each of the endogenously tagged Nups of interest and to characterize the so far unmapped position of ELYS.

**Figure 1.**
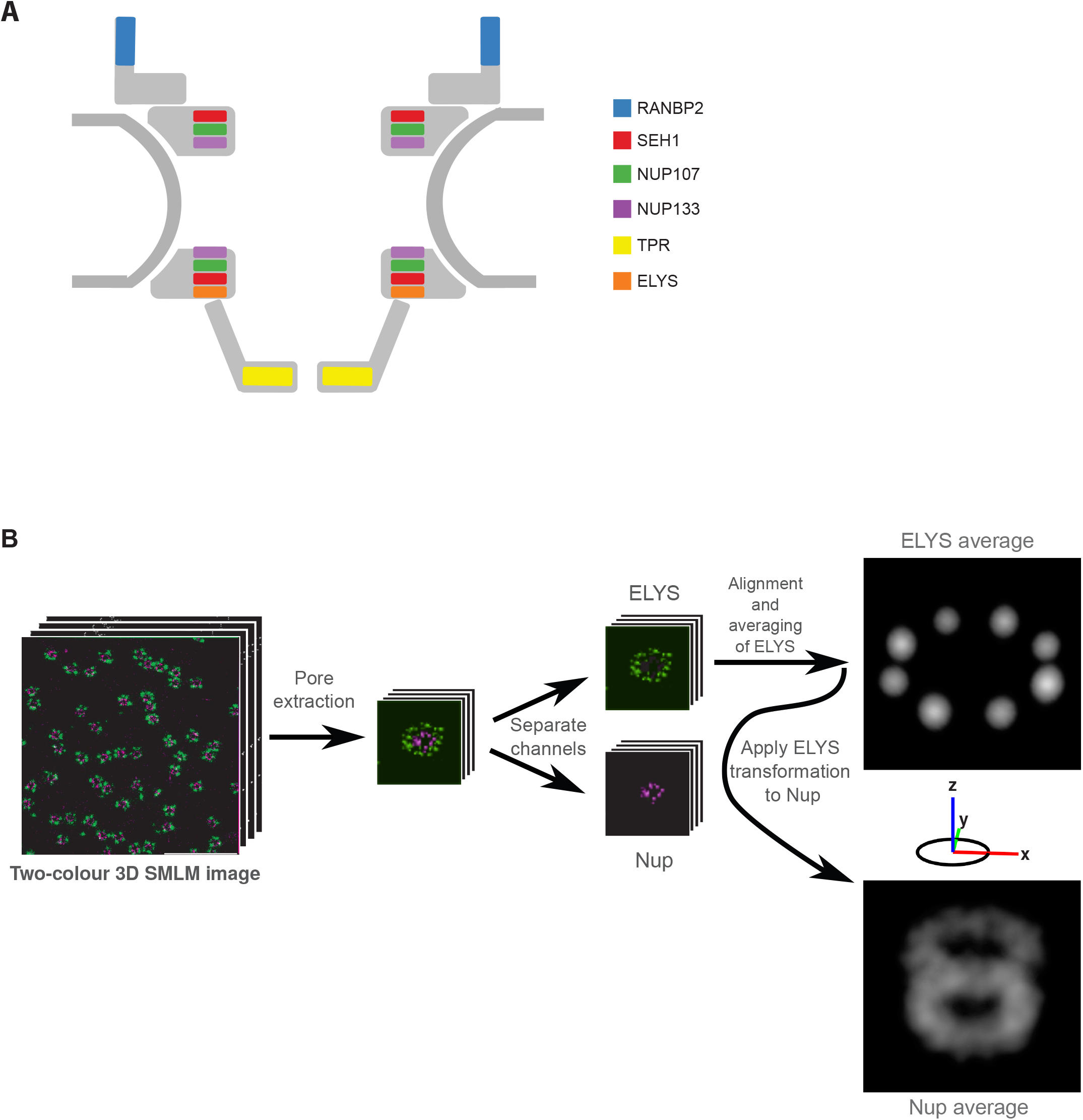
3D averaging of SMLM images of the nuclear pore complex. **A**) Schematic representation of the NPC indicating the position of the proteins imaged in this study. **B**) Schematic representation of the 3D SMLM averaging pipeline. From a dual-colour 3D SMLM image stack of the lower surface of the nucleus of a U2OS cell, endogenously expressing a SNAP-Nup labeled with BG-AF647 and ELYS visualized with antibodies coupled to CF680, individual pores were automatically picked and an iterative 3D alignment process was applied to the ELYS channel imposing an 8-fold rotational symmetry. The same transformation was then applied to register the Nup of interest in the other channel to preserve the spatial relationship between the two labelled proteins. The final reference image was visualized as an intensity-rendered 3D reconstruction.

To analyze the 3D single molecule localizations obtained from many NPCs, we further developed a computational pipeline initially used for subtomogram averaging (Beck et al., 2004) that integrates the information of thousands of 3D SMLM sub-volumes containing individual NPCs and extracts quantitative 3D information of the average distribution of each Nup. This pipeline consists of three major steps: (i) automatic sub-volume extraction of single NPCs, (ii) iterative generation of reference and (iii) iterative sub-volume alignment. We used this pipeline to analyze the ELYS reference channel imposing the well-known 8-fold symmetry of the NPC in the averaging and then applied the same transformations to the channel of the Nup of interest. The final averaged reference is then visualized as a 3D projection of a localization-density map of both channels. It recovers the precise 3D position of the ELYS reference and contains quantitative information about the averaged axial and radial positions of the Nup of interest relative to ELYS (**Fig. 1B**; for details see Methods).

The vertebrate Nup ELYS has been characterized as a member of the scaffold of the NPC (Franz et al., 2007), however available cryo-EM density maps could not capture its exact position (von Appen et al., 2015), and thus suggested its localization must be asymmetric within the NPC. Our mapping of ELYS by super-resolution microscopy revealed a single unexpectedly large ring (diameter: 140 nm) (**Fig. 1B)**. Comparison to the position of the scaffold Nups SEH1 and NUP107 allowed us to place it underneath and outside the nuclear ring **(Fig. 2B–C)**, providing the first direct evidence that it is indeed asymmetrically positioned on the nucleoplasmic side of the NPC (**Fig. 2A–E)**, consistent with previous computational models (von Appen et al., 2015).

**Figure 2.**
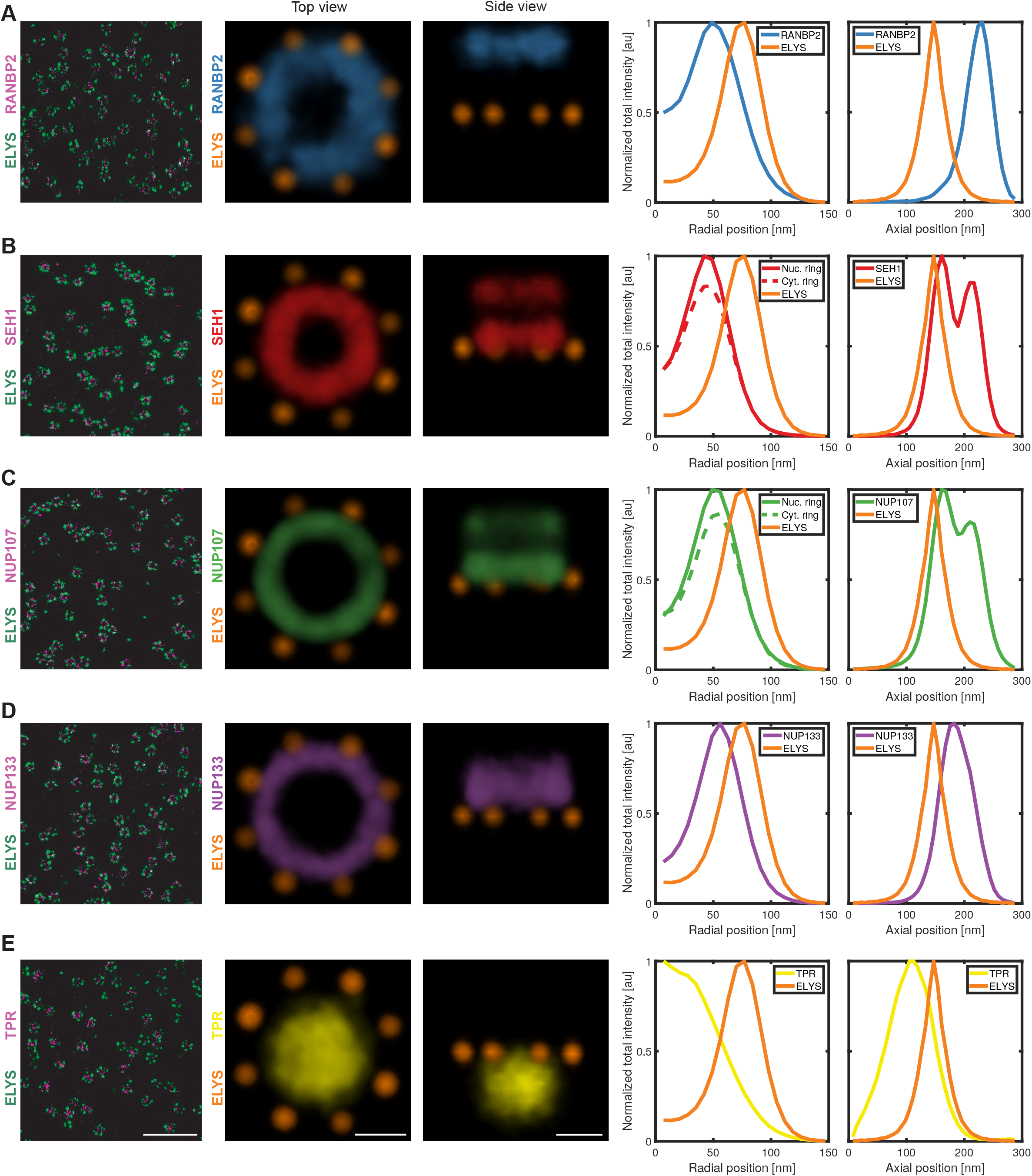
Average intensity map of individual Nups. Representative dual-color 9-μm^2^ area of the lower surface of the nucleus of a U2OS cell, endogenously expressing **A**) SNAP-RANBP2, **B**) SNAP-SEH1, **C**) SNAP-NUP107, **D**) SNAP-NUP133, **E**) TPR-SNAP, respectively, labeled with BG-AF647 and with α-ELYS primary antibody and secondary coupled to CF680. Top-view and side-view of density maps of the 3D averages of 3D SMLM datasets (RANBP2 (n = 2140): light blue, SEH1 (n = 1619): red, NUP107 (n = 3617): light green; NUP133 (n = 2418): purple, TPR (n = 3066): yellow and ELYS: orange). Scale bars, 1 μm for overview images and 50 nm for average intensity map.

Relative to ELYS, we then examined the structure of three members of the largest NPC subcomplex, the Y-shaped complex, SEH1, NUP107 and NUP133, that are expected to be present in two symmetric, nuclear and cytoplasmic rings. Our 3D averaged maps based on the ELYS reference indeed recovered this double ring architecture for SEH1 and NUP107, while only a single thicker ring could be resolved for NUP133, presumably due to them being stacked most closely together (**Fig. 2B–D**). SEH1 had the smallest ring diameter (84 nm) and the largest separation between the nucleoplasmic and cytoplasmic ring (49 nm), with its nucleoplasmic ring only 14 nm away from ELYS (**Fig. 2B**, **Table 1**). Slightly more central then followed the larger NUP107 double ring (diameter: 98 nm, axial distance: 43 nm, distance to ELYS: 17 nm). The ELYS based average distribution of NUP133 did not allow us to clearly resolve a double ring, but rather an even larger (103 nm diameter) and thicker ring, positioned more centrally and further away from ELYS (38 nm) **(Fig. 2D and Table 1)**. The ring diameters of the three scaffold Nups are consistent with our previous mapping of their radial positions by 2D super-resolution microscopy (Szymborska et al., 2013), and we now provide direct evidence for their axial positions. Our data is furthermore consistent with current models of NPC structure based on in vitro data (von Appen et al., 2015), additionally positioning and providing novel evidence of the asymmetric nature of ELYS; validating our methodological pipeline for mapping the 3D molecular architecture of multiple nuclear pore components by fluorescence microscopy at the nanoscale in situ.

**Table 1.** Quantitative analysis of the architecture of the NPC. The diameter, distance between rings, distance to the reference (ELYS) and spread along z axis and xy plane are calculated from the corresponding dual-color 3D averaged images.

|  | RANBP2 | SEH1 | NUP107 | NUP133 | TPR | ELYS |
| --- | --- | --- | --- | --- | --- | --- |
| **Diameter** | 95 nm | 84 nm | 98 nm | 103 nm | NA | 137 nm |
| **Distance between rings** | NA | 49 nm | 43 nm | NA | NA | NA |
| **Distance from ELYS** | 80 nm | 14 nm | 17 nm | 38 nm | 40 nm | 0 |
| **Spread along z axis** | 52 nm | 95 nm | 97 nm | 71 nm | 92 nm | 42 nm |
| **Spread along xy** | NA | NA | NA | NA | 100 nm | NA |

The cytoplasmic filaments and the nuclear basket are the least structurally characterized components of the NPC. Interestingly, 3D dual-color reconstructions of these subcomplexes revealed for RANBP2, a major component of the cytoplasmic filaments, a single ring of 100 nm in diameter, displaced 80 nm towards the cytoplasm from the ELYS reference ring (**Fig. 2A; Table 1**). Images of the nuclear basket protein TPR showed an unexpectedly large, slightly flattened ellipsoid shaped volume with 100 nm in diameter at its central base and 92 nm diameter at its distal cap, displaced 40 nm towards the nucleoplasm from the ELYS reference ring, capturing for the first time the 3D distribution of the major component of the nuclear basket (**Fig. 2E; Table 1**).

Next, we integrated all our data into a unified 3D multimolecular map that reveals the positions of all investigated Nups relative to each other **(Fig 3**). The constant reference ELYS allowed the interpolation and alignment of all the dual-color reconstructions into a single multicolor map (**Fig. 3A**). This map provides a representation of the average positions of all imaged Nups and allows a comprehensive quantitative analysis between both symmetrically and asymmetrically positioned components (**Fig. 3C**). Our integrated 3D map of the NPC revealed, for example, that RANBP2 is localized exclusively at the cytoplasmic ring more peripheral but with a slightly larger diameter compared to the most cytoplasmic part of the scaffold formed by SEH1 (**Fig. 3A–B**). Furthermore, the 3D map showed that ELYS resides exclusively in the nucleoplasmic ring with a much larger diameter and more peripherally than the most nuclear part of the scaffold, also formed by SEH1. Maybe most strikingly, the nuclear basket filament protein TPR is distributed over a large volume, reaching much further towards the nucleoplasm than ELYS, on average appearing to explore the whole opening of the stacked ring channel formed by all other NPC components if viewed along the transport axis (**Fig. 3B**).

**Figure 3.**
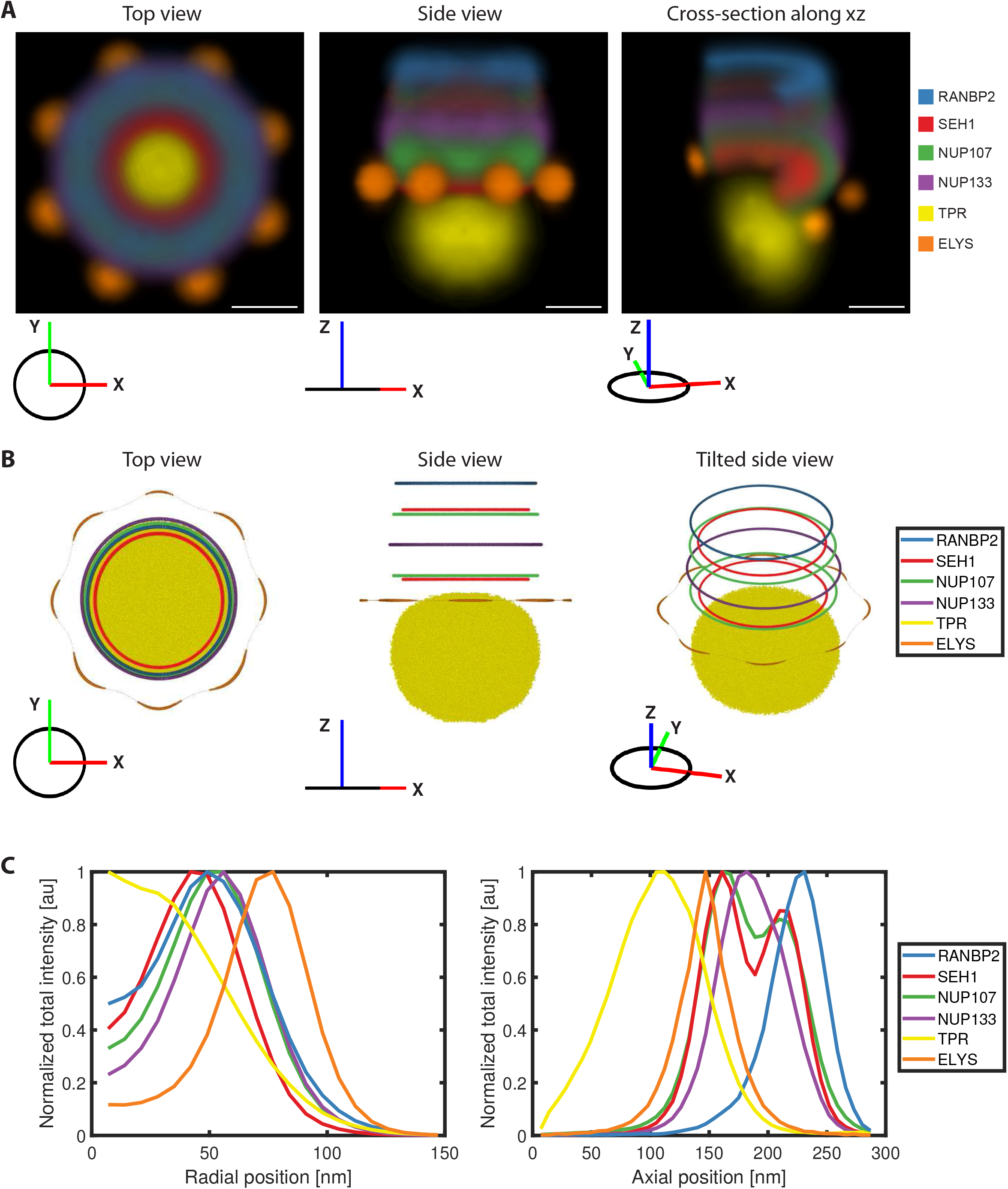
Integrative 3D map of the NPC. (**A**) Density map of the top (left), side (middle) and tilted side-view of cross-section along the x-y plane (right) of the combined average dual-color 3D SMLM datasets (RANBP2: light blue, SEH1: red, NUP107: light green; NUP133: purple, TPR: yellow and ELYS: orange). Rotational averaging around the *z*-axis was performed on individual average intensity maps to generate integrative 3D map. The orientation of the density map is indicated as a coordinate system for each view. Scale bar, 50 nm. (**B**) Center of the rings of different Nups calculated from the integrated average maps in (A) and intensity rendering of top 45% density of TPR. (**C**) Radial (left: indicating intensity distribution from center to the periphery) and axial (right: indicating intensity distribution along z-axis) total intensity profiles of different Nups, normalized to the maximum intensity of individual profiles.

Similar to most existing NPC reconstructions based on other approaches, our integrated 3D model represents an average structure of the pore and thus assumes a single consensus conformation (von Appen et al., 2015; Bui et al., 2013; Kim et al., 2018; Kosinski et al., 2016; Lin et al., 2016). By construction, it therefore dismisses information about possible structural variability within the NPC. However, the unexpectedly large volume filled in our average map by combining all localizations we obtained for TPR, could for example result from a combination of different conformations. Furthermore, the fact that we did not recover the 8-fold subunit arrangement of the individual Nups when using the 8-fold symmetry we assumed for ELYS as reference, would also be consistent with structural variability between different NPC substructures. Since fluorescence microscopy can obtain a comparatively high signal-to-noise ratio for individual NPCs in SMLM sub-volumes and has the a priori molecular identity of each labeled protein, we wanted to explore if our data allows us to go beyond averaging of all NPCs and probe the presence of different structural states that might be expected in the natural environment of the cell. To complement our particle averaging approach, we therefore decided to quantitatively analyze the variability among single NPCs in our 3D super-resolution images.

To initially explore the plasticity of the NPC for each of the thousands of individual 3D SMLM sub-volumes (**Fig. S1A)**, we first quantified the degree of heterogeneity for simple geometric parameters of the rings formed by scaffold Nups and RANBP2, measuring their diameter, roundness, spread along *z*-axis and distance between two rings, where present. This analysis revealed that, while the median values are consistent with those measured from the reconstruction (**Table 1**), there is also significant variation in all parameters, such as circularity and diameter of the different ring-like structures (**Fig. S1B, C**). Furthermore, the center of mass along the transport axis, as well as the distance between the double rings formed by SEH1 or NUP107 along the transport axis also varied substantially (**Fig. S1D, E)**. It is important to note that the localization precision of ~7 nm in our 3D SMLM experiments, which we determined as described previously (Ries, 2020; Li et al., 2018), and the size of the labels (antibodies ~15 nm, SNAPtag ~2 nm) leads to a random spread of the localizations around the true protein positions. In addition, systematic errors due to sample induced aberrations (Li et al., 2018) and residual registration errors further add to the technical variability. However, the heterogeneity we detect explores a larger range of distances, indicating that it also reflects variable structural properties of the NPC. Taken together, the exploration of single NPC derived geometric parameters was consistent with the presence of different structural conformations for both scaffold and peripheral Nups in our data set.

To assess if the NPC is indeed present in multiple conformational states, we explored our data with an orthogonal approach that compares all single molecule localizations obtained for each NPC sub-volume in an unbiased manner. To this end, we developed an unsupervised classification methodology based on topological data analysis (TDA) using persistent homology based clustering (Edelsbrunner et al., 2002), which allows the robust classification of 3D localization point clouds based on topological structure similarity alone, without assuming any structure or symmetry a priori (Hofmann et al., 2018). In brief, each labeled instance of a nuclear pore is first represented by the point set of its fluorophore positions after which a topological signature is derived by connecting all fluorophore positions of the pore within distance *t* of each other and analyzing the connectivity of the whole point set for a range of values of *t*. Using a similarity measure between the topological signatures allows clustering the pores into structurally similar classes. Because this approach only relies on pairwise distances between points, the resulting signature captures a notion of shape beyond the presence of ring structures that is structurally unbiased and for example independent of position and rotation. It is expected that both the size and shape of the rings would also be captured by the topological signatures and therefore influence the clustering. While this structurally unbiased classification is powerful, its limitation is that it does not allow an a priori interpretation of the clusters in terms of geometric features.

To validate our classification methodology, we first tested if it could distinguish the different imaged Nups from each other based on topological features alone. To this end, we applied hierarchical clustering to the topological signatures of the whole data set of more than ten thousands NPCs **(Fig. 4A)**. The first split in the tree separates most of the Y-complex Nups, i.e. NUP107, NUP133 and SEH1, from more peripheral Nups RANBP2 and TPR and despite some noise, pores labeled for the same Nup tend to cluster together. To quantify the performance of the approach, the tree was cut to generate subclusters. The cut height was chosen to allow capture of the smaller clusters resulting from underrepresentation of some of the proteins in the data. Each of the 22 clusters was dominated by one Nup with average cluster purity of 66% (ranging from 33% to 100%). To further evaluate how these clusters capture the underlying ground truth represented by the known Nup identity, we built the confusion matrix resulting from assigning the dominant Nup as cluster identity (**Fig. 4B**). This analysis shows that this simple classification scheme performs moderately well with an overall accuracy of 53% which is significantly above the no-information rate (p-value < 2.2e^**−**16^). The least well predicted Nup is SEH1, which is most often misclassified as NUP133. The fact that even subsets of Nups that showed a very similar double ring structure such as SEH1 and NUP107 could be distinguished in this manner, illustrates that persistent homology-based clustering is a robust unbiased method for the classification of 3D SMLM images.

**Figure 4.**
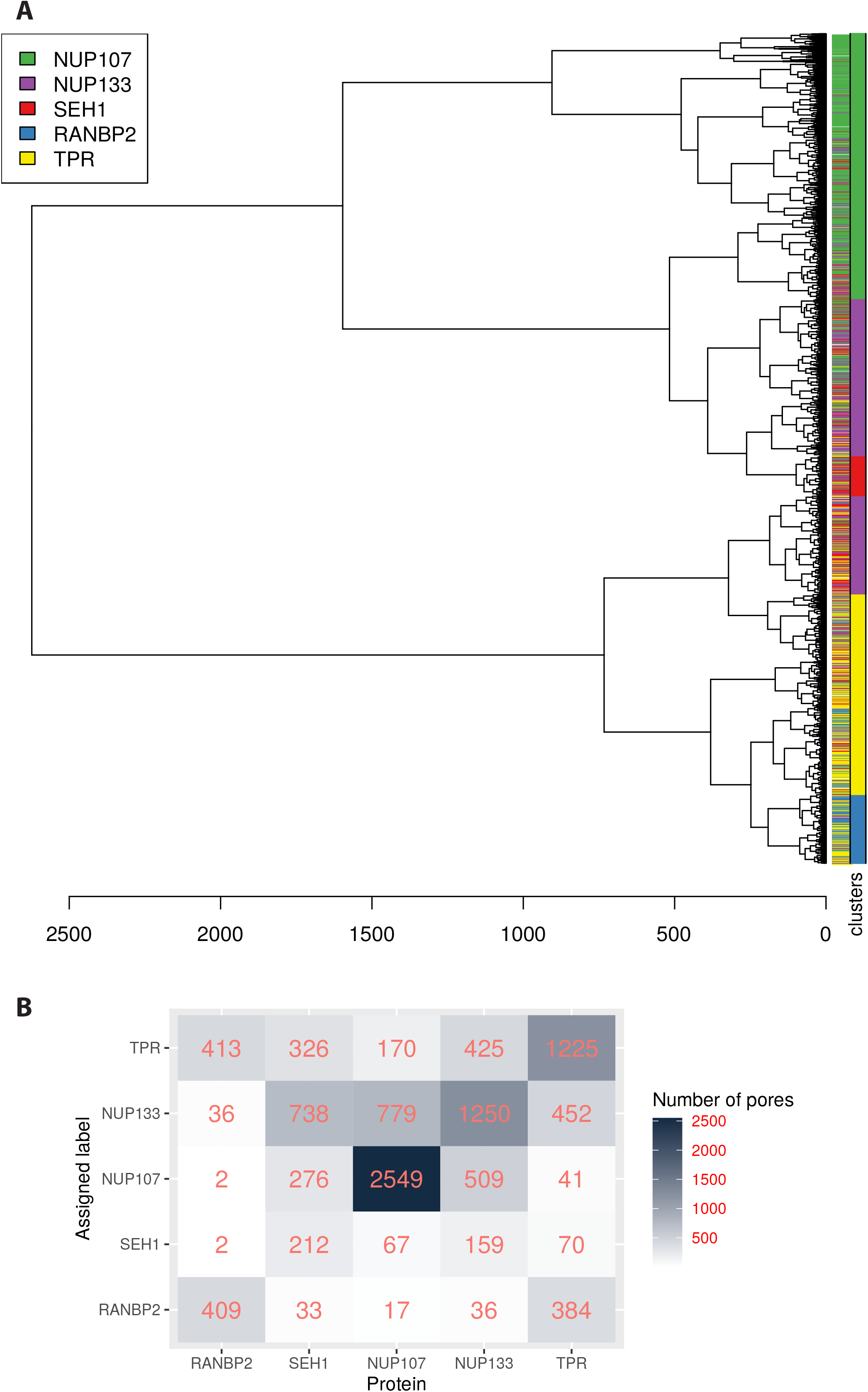
Clustering of persistence diagrams of Nups of interest. (**A**) Dendrogram of the persistent homology clustering of 10580 3D SMLM particles representing all the Nups of interest. Clusters were identified by cutting the dendrogram at a height of 200 (red line) and each cluster was assigned the identity of the dominant Nup represented in it (coloured bar). Although particles for each Nup tend to group together, there is some overlap with other Nups with similar structure and the existence of distinct clusters for each Nup suggests the existence of structural diversity of the NPC (SEH1: purple; NUP107: pink; NUP133: green, RANBP2: blue and TPR: yellow). See Materials and Methods for details on clustering. (**B**) Confusion matrix showing the classification outcome of the clustering scheme in (A). The mean balanced accuracy is 68%.

Next, we investigated if topological data analysis can also identify different conformational states between the individual NPCs imaged for one specific Nup. Using the branching inferred from the global similarity tree for one protein at a time, classes were defined by cutting the tree to produce four to five groups containing a sufficient number of NPCs to be analyzed statistically **(Fig. 5)**. To explore if the different classes represent different structural conformations, we used the two approaches we had already successfully used for the global average structural analysis, particle averaging and geometric parameter analysis. First, we separately 3D averaged the particles of one Nup within each class. Two classes were identified for RANBP2. These revealed differences in the extent of the central opening (**Fig. 5A, B**) suggesting that RANBP2 is not strictly confined to a ring structure. The scaffold Nups SEH1 and NUP107, where double rings could be detected, clustered into four to five classes. Particle averaging of each class revealed one major architectural feature distinguishing them. Intriguingly, we found either a completely open, or a partially or almost fully filled nuclear ring structure (**Fig. 5 C,D, clusters 3 and 4; E,F, cluster 4**). The classes with filled ring structures had fewer members, but still represented over 100 NPCs, each of which exhibited comparable numbers of localizations to NPCs in other classes, indicating that they represent a different structural state, rather than poorly labeled particles. Our unsupervised classification by topological data analysis thus confirmed the presence of consistently different conformational states among NPCs. Very interestingly, the 5-10% of filled ring structures we observed for scaffold Nups is consistent with the fraction of assembling pores previously estimated to be present in growing interphase nuclei (Dultz et al., 2010; Otsuka et al., 2016) and it is therefore tempting to speculate that they might represent NPC assembly intermediates.

**Figure 5.**
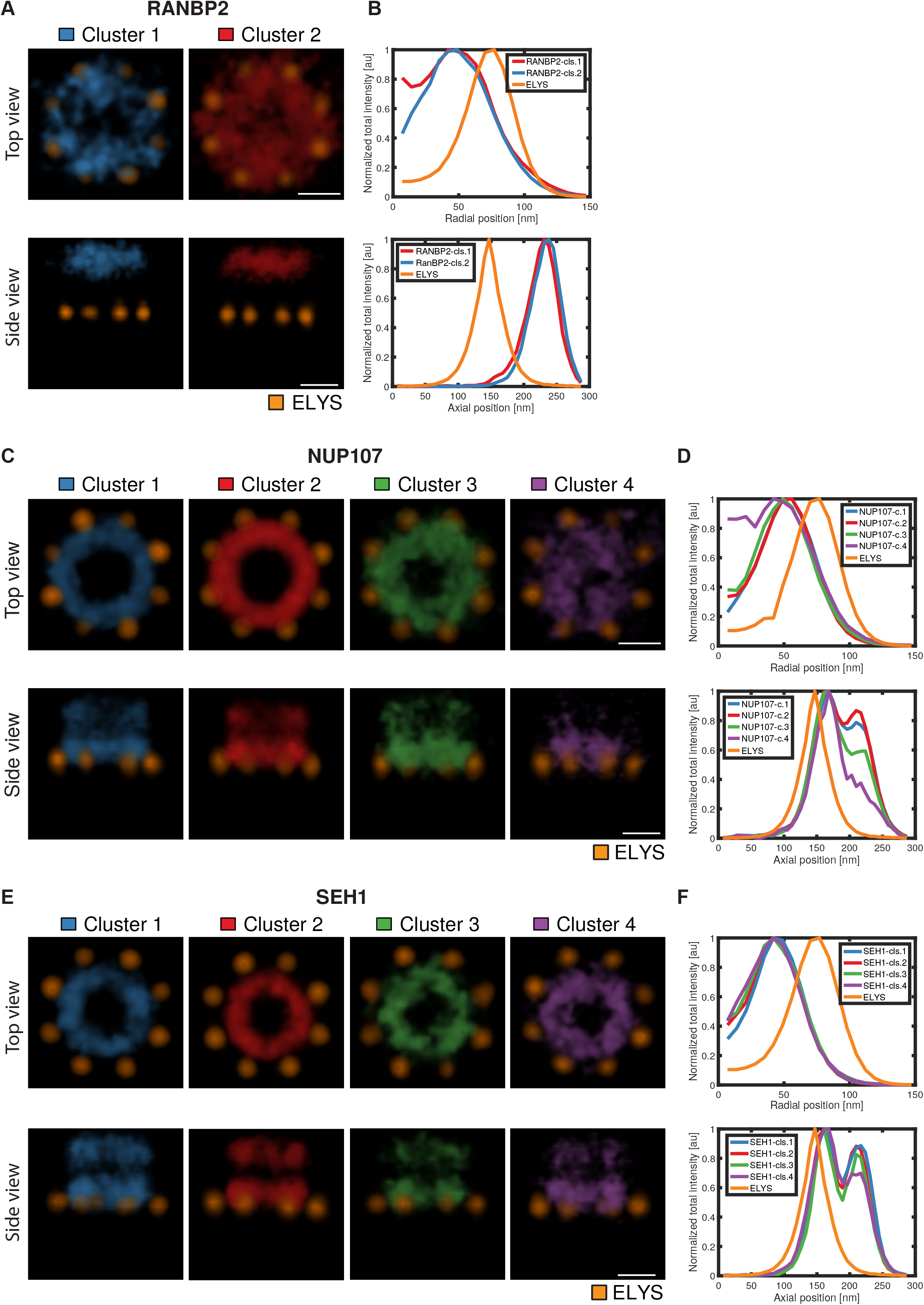
Conformational states of the NPC. (**A**) Volume-rendered density map of RANBP2 clusters. Scale bar, 50 nm. (**B**) Radial and axial distributions of signal for each RANBP2 cluster. (**C)** Volume-rendered density map of NUP107 clusters. (**D**) Radial and axial distributions of signal for each NUP107 cluster. Scale bar, 50 nm. (**E**) Volume-rendered density map of SEH1 clusters. (**F**) Radial and axial distributions of signal for each SEH1 cluster. Scale bar, 50 nm.

## Discussion

This work combines 3D super-resolution microscopy of pairwise combinations of knocked-in Nups with a constant reference in human cells with particle averaging to create an integrative 3D multi-molecular map of the NPC. Our average map is in good agreement with the currently available high-resolution cryo-EM structural model of the symmetric scaffold of the human NPC (von Appen and Beck, 2015; Bui et al., 2013; Kosinski et al., 2016) and provides the first direct evidence for several aspects of the model that were so far only predicted. Importantly, our map includes for the first time also asymmetrically positioned Nups that were absent or ill-defined in the cryo-EM model, thereby providing novel evidence of the molecular position of the scaffold nucleoporin ELYS, the cytoplasmic filament component RANBP2 and the core element that forms the nuclear basket, TPR. Our data for example revealed that the ring structure formed by eight ELYS containing subunits exhibits a significantly larger diameter than even the most peripheral scaffold NUP107, even when taking into account the potentially larger fluorophore displacement of an antibody (Ries et al., 2012) compared to the SNAP tag. ELYS has been reported to bind to chromatin (Rasala et al., 2006) and the position we found for it outside of the scaffold rings and away from the large central volume explored by the nuclear filament protein TPR could allow ELYS to stably anchor the core NPC to chromatin without interfering with the transport process.

Beyond these new insights from a multimolecular structure model that includes so far unmapped components, our unsupervised computational classification of all individual NPCs revealed the presence of different structural classes of NPCs for specific Nups, highlighting a small fraction of potential assembly intermediates with filled in nuclear rings. Why did previous structural studies of the NPC not notice this? In the main approach taken so far, cryo-electron tomography with subtomogram averaging, very flexible structures such as the nuclear basket are completely absent. Even for the more stable core scaffold however, the most prominent conformational state will typically dominate the averaging and hide a small fraction of different conformers. An in-depth classification of very large numbers of relatively low contrast cryo-EM sub-volumes that would be needed to detect assembly intermediates has not been carried out so far. However, alternative approaches by scanning electron microscopy have suggested the presence of partially assembled or “mini” pores in the past (Maeshima et al., 2010 Nat Struc Biol), which is consistent with our super-resolution microscopy-based observations reported here.

Overall, our approach of mapping the multimolecular architecture of large protein complexes by super-resolution microscopy and computational analysis constitutes an important complementary methodology to higher-resolution methods that will allow the investigation of the structural diversity and function of molecular machines in their natural cellular environment. In the future, this approach will furthermore allow for directly linking different functional states or perturbations to different molecular structures in single cells.

## Contributions

V.J.S and J.E conceived the project, A.S., J.R and M.B contributed to the conception and design of the work, B.N., M.K., A.C and V.J.S. generated and validated the cell lines, V.J.S. and B.N performed the experiments, V.J.S., and P.H., acquired and processed the data, V.J.S, J.K.H, M.J.H., S.M., P.H., J.R., M.B., developed the methods, wrote the software and analyzed the data. V.J.S and J.E wrote the manuscript. All authors contributed to the interpretation of the data, read and approved the final manuscript.

## Acknowledgements

We thank S. Correia, N. Daigle and B. Koch for experimental support; A. Rybina, Otsuka and A. Politi for discussion and S. Alexander for critical reading of the manuscript.

This work was supported by the Baden-Württemberg Stiftung (to J.E.), the European Research Council (grant no. ERC CoG-724489 to J.R.), the National Institutes of Health Common Fund 4D Nucleome Program (grant no. U01 EB021223 to J.R. and J.E.), the Human Frontier Science Program (grant no. RGY0065/2017 to J.R.) and the European Molecular Biology Laboratory (EMBL; all authors). V.J.S. acknowledges support by the Boehringer Ingelheim Fonds. P.H. is a candidate for joint PhD degrees from EMBL and Heidelberg University.

## Materials and Methods

### Cell culture

The human osteosarcoma U2OS (RRID: CVCL_0042) cells were purchased from ATCC (Wesel, Germany). U2OS cells were cultured 1x high glucose McCoy’s 5A modified medium (McCoy; Thermo Fisher Scientific, cat.# 26600023). The McCoy media was supplemented with 10% (v/v) fetal bovine serum (FBS; Thermo Fisher Scientific; cat.#10270106; qualified, E.U.-approved, South America origin), 100 U/ml penicillin-streptomycin (Thermo Fisher Scientific; cat.# 15140122), 2 mM L-glutamine (Thermo Fisher Scientific; cat.# 25030081), 1 mM sodium pyruvate (Thermo Fisher Scientific; cat.# 11360070) and 1% MEM nonessential amino acids (Thermo Fisher Scientific; cat.# 11140050). Cells were grown on culture dishes (ThermoFisher) at 37 °C and 5% CO2 in a cell culture incubator. Cells were passaged at a confluence of around 80% every 3 days by trypsinization (Trypsin-ethylenediaminetetraacetic acid (EDTA; 0.05%), phenol red (Thermo Fisher Scientific; cat.# 25300054)). Tests confirming the absence of mycoplasma were performed every 2 months.

### Cell line generation and validation

Genome editing was carried out in U2OS cells in order to express endogenously tagged nucleoporins of interest (NOIs). For tagging SEH1, NUP133, NUP358/RANBP2 at the N-terminus and TPR at the C-terminus with SNAPf-tag paired CRISPR/Cas9D10A nickases were used. For tagging NUP107 at the N-terminus with SNAPf-tag Zinc finger nucleases (ZFNs) were used. The design, cloning and transfection were carried out as described in (Koch et al., 2018; Mahen et al., 2014). After the first round of genome-editing homozygous clones were achieved from NUP107, SEH1, NUP133 and NUP358/RANBP2 and heterozygous clones for TPR tagged expression. A second round of genome editing was performed on the heterozygously-tagged TPR expressing clones after confirming with Sanger sequencing the absence of mutations in the wild-type alleles. All endogenously tagged cell lines were validated by our standard validation pipeline described in (Koch et al., 2018).

### Sample preparation

The homozygously expressing tagged-NOIs U2OS were seeded on 24 mm #1.5 round coverslips for 3D SMLM, which had been washed overnight in a 1:1 mixture of methanol and hydrochloric acid, rinsed with dH_2_O until neutral and subsequently sterilized with UV light. Seeded cells were allowed to attach overnight at 37°C and 5% CO_2_ in a cell culture incubator. The coverslips containing the homozygously expressing SNAP-NOIs cells were pre-fixed with 2.4% [w/v] formaldehyde (FA; Electron Microscopy Sciences; cat.# 15710) in 1x PBS (+Ca^2+^/Mg^2+^), permeabilized with 0.5% [v/v] Triton X-100 (Sigma-Aldrich; cat.# T8787) in 1x PBS (+Ca2+/Mg2+), fixed afterwards with 2.4% FA and quenched for remaining fixative with 50 mM NH_4_Cl. The fixed samples were blocked with a few drops of Image-iT FX Signal Enhancer (Thermo-Fisher; cat.# I36933). The benzylguanine (BG)-conjugated AF647 (New England Biolabs, cat.# S9136S) was diluted to 1 μM in blocking solution (0.5% (w/v) BSA, 1 mM DTT in 1x PBS) and incubated with the sample. We note that labelling efficiency of double ring Nups appears asymmetric with the cytoplasmic ring less efficiently labelled than the nuclear ring. The samples were blocked again with 5% Normal Goat Serum (NGS; Sigma-Aldrich; cat.# G9023) in 1x PBS (+Ca^2+^/Mg^2+^). The primary anti-ELYS antibody (polyclonal rabbit anti-AHCTF1-antibody; Sigma-Aldrich; cat.# HPA031658; 1:30) was diluted in blocking solution (5% (w/v) NGS in 1x PBS) and incubated with the sample. This antibody was produced and validated by the Human Protein Atlas project and is detailed at this URL: https://www.proteinatlas.org/ENSG00000153207-AHCTF1/antibody#antigen_information. The antigen is located towards the C-terminus of the protein, between two domains, the ELYS domain and the C-terminal AT-hook domain in a region with no known or detectable structure.

The secondary antibody coupled to CF680 (Sigma; cat.# SAB4600362 1:1000) was diluted in blocking solution (5% (w/v) NGS in 1x PBS) and incubated with the sample.

### 3D SMLM microscope and imaging

All 3D SMLM data were acquired on a custom-built wide-field setup described previously (Deschamps et al., 2016) using a cylindrical lens (f = 1000 mm; Cat#LJ1516L1-A, Thorlabs) to introduce astigmatism. First, *z*-stacks with known displacement of several (15-20) fields of view of TetraSpeck beads on a coverslip were acquired to generate a model of the experimental point spread function (Li et al., 2018). This model was then used to determine the *z*-position of the individual localizations. Emitted fluorescence was collected through a high-numerical-aperture (NA) oil-immersion objective (160x/1.43-NA; Leica, Wetzlar, Germany), filtered by a bandpass filter 700/100 [Cat#ET700/100m, Chroma] for AF647 and CF680) and imaged onto an Evolve512D EMCCD camera (Photometrics, Tucson, AZ, USA). For ratiometric dual-color imaging of AF647 and CF680, the emitted fluorescence was split by a 665LP beamsplitter (Cat#ET665lp, Chroma), filtered by a 685/70 (Cat#ET685/70m, Chroma) bandpass filter (transmitted light) or a 676/37 (Cat#FF01-676/37-25, Semrock) bandpass filter (reflected light) and imaged side by side on the EMCCD camera. The color of the individual blinks was assigned by calculating the ratio of the intensities in the two channels.

Coverslips containing prepared samples were placed into a custom built sample holder and 500 μL of GLOX-buffer (50 mM Tris (Sigma-Aldrich; cat.# T1503), pH 8; 10 mM NaCl (Merck; cat.# 106404); 10% (w/v) D-glucose (Merck; cat.# 104074), 0.5 mg/mL glucose oxidase (Sigma-Aldrich; cat.# 49180); 40 μg/mL catalase (Sigma-Aldrich; cat.# C3155); and 35 mM MEA (Sigma-Aldrich, cat.# 30070). The buffer solution was exchanged after about 2 h of imaging. Samples were imaged until close to all fluorophores were bleached and no further localizations were detected under continuous UV irradiation.

### 3D SMLM data analysis

All data analysis was performed with custom software written in MATLAB and is available as open source (Ries, 2020). All 3D SMLM data were fitted and analyzed as described previously (Li et al., 2018). The 3D SMLM images were rendered as localization histograms with a bin size chosen to be 7 nm and exported as TIFF stacks for further analysis.

### Image analysis

#### Software and Toolboxes

All analysis was performed using MATLAB R2017b, R2018 and R2019 (Mathworks) and R versions 3.4.2 – 3.6.1 (R Foundation for Statistical Computing, 2018). The averaging operations were carried out with TOM (Nickell et al., 2005) and av3 (Forster et al., 2005) toolboxes in MATLAB.

#### NPC particle picking

The individual NPC particles were selected from the 3D SMLM image-stacks by an automatic particle-picking pipeline developed in the R programming language (R Foundation for Statistical Computing, 2018). The automatic particle-picking pipeline implements two different algorithms for identifying NPC particles: a density-based clustering algorithm (DBSCAN) (Ester et al., 1996) (and a template-matching algorithm (circle Hough transform) (Duda and Hart, 1972). After detecting all possible NPCs within an image-stack, the selected particles are cropped into image-stacks of 40×40×40 voxels, where each voxel corresponds to 7 nm.

#### 3D SMLM averaging

To keep the spatial relationship between ELYS and Nups, averaging is performed on signal from the ELYS channel and the same transformation is applied to register the other channel with the Nup of interest. An initial ELYS reference volume was generated by the sum of all automatically cropped particles. An iterative averaging algorithm was used for aligning the particles and to refine the initial reference (Beck et al., 2004). Briefly, the alignment was performed by angular search of the 3D SMLM particles (i.e. both rotational and translational offset are calculated by 3D cross correlation (CC) in Fourier space), to generate the subsequent references. Here, the orientation of each particle is described by the Euler angles φ, ψ, and θ. An initial averaging model of the NOIs was generated using a random distribution of the polar angle φ. During the iterative refinement, eight-fold symmetry along φ was imposed (i.e. based on the known NPC symmetry) in order to improve the SNR. Ideally, this algorithm takes into account the ‘missing wedge’ present for each of the sub volumes (tomograms) (Beck et al., 2004; Nickell et al., 2005) and given that 3D SMLM data contains no ‘missing wedge’, this parameter was supplemented with full angular coverage of −90° to 90°. The averaging was run for ten iteration cycles for all Nups of interest and their clusters and it was observed that ten iterations were sufficient for angular convergence. Once averaging of ELYS was performed, the same transformations of individual particles were applied to register the associated Nups. Averaged ELYS from different Nups were further registered to bring all the Nups of interest into the same coordinate system which allowed visualization of integrated maps and made extraction of different parameters possible **(Fig. 3 and Table 1)**. For visualization purposes and generation of the photon density maps the final reconstructions were interactively visualized and analyzed in the molecular modeling software UCSF Chimera (Pettersen et al., 2004).

#### Extraction of different parameters to analyse the position and distributions of Nups

The axial and lateral positions of the 3D SMLM averaged 3D reconstructions were calculated by fitting a Gaussian function to their average profiles projected in different ways **(Fig. S2)**. To estimate radial intensity distributions, radial profiles were created from the center of the image to the periphery at 1-degree angular rotation with respect to the z-axis and summing up the intensities in different z-slices along the profile. The average radial profile was generated by summing up all 360 profiles. **Fig. S2A** shows construction of two profiles taken at 0 and 30 degrees. The center of the ring defining the radius was calculated by fitting a Gaussian to individual radial profiles. The diameter is calculated by taking the average of all radii multiplied by two. The axial profile was generated by the sum projection along the *z-*axis of the average image. The spread of Nups along the z-axis was calculated by taking the full width at half maximum (FWHM) on this profile **(Fig. S2B)**. The spread in XY was calculated by taking the mean of spreads along x and y axes. To get the distance between rings for Nups forming double rings (NUP107 and SEH1), parts of stack containing nuclear and cytoplasmic rings were analyzed separately where the separation point was decided from the valley of their axial intensity profile. An axial intensity profile for each of these coordinates was created by taking the total localization signal in a window of 3×3 pixels centered at the coordinate in each slice within the *z*-stack. A Gaussian function was fitted to this profile and the axial position of a ring was defined by the mean of all fitted positions.

#### Quantifying diameter and height in individual pores

Localization signals of individual registered NPC particles were sum projected along the *z*-axis. Projected images were then transformed to polar images of size 21 × 360 pixels where the *x*-axis represents the angles in [1, 360^0^] and *y*-axis represents corresponding intensity profiles of the Cartesian image. Lateral (radial) distances along 4 angular directions (0, 90, 180 and 270 degrees) were computed by fitting a Gaussian function to the profiles constructed from summing up 90 columns in the polar image starting from 1 to 90 and ending at 271 to 360 degrees. The diameter of an NPC particle was defined as two times the mean of lateral distances along all angular directions where some localization signal is available. The distance between two rings was calculated in the same way as calculated for average Nups, but measurements were taken only at 4 angular positions.

#### Quantifying roundness of individual pores

Considering the fact that individual particles have a smaller number of positive voxels, the roundness of a particle was measured by the ratio of two eigenvalues computed from 2D projected coordinates of its localization signals. Coordinates of all the voxels having some localization signals were used to construct a Hessian matrix. Three orthogonal eigenvectors and three associated eigenvalues were calculated from this matrix. The roundness of the ring was defined by the ratio between the square roots of the second to the first eigenvalues of this matrix.

#### Quantification of spread along z direction

Total signal in each plane of the *z*-stack was computed which resulted in a one-dimensional vector of size 41. The spread of Nups along z-axis was calculated by taking full width at half maximum (FWHM) on this profile **(Fig. S2B)**.

#### Topological data analysis

Each NPC was represented by a point set defined by the 3D Cartesian coordinates of the localization signals. To characterize differences in conformations of structures represented by these 3D point sets, we turned to topological data analysis. For the purpose of topological data analysis, data is viewed as a topological space from which information about the structure of the data can be inferred. In practice, this means considering a set of data points with a measure of distance between them as representing an underlying topological space and extracting relevant topological characteristics. A powerful tool to derive features from such data is persistent homology (Edelsbrunner and Harrer, 2010) where the *k*-dimensional homology group characterizes the topological features seen in that dimension, i.e. 0-dimensional homology characterizes connected components, 1-dimensional homology characterizes loops (or equivalently holes inside loops) and 2-dimensional homology characterizes voids (i.e. bubbles). Persistent homology efficiently computes homology groups at multiple scales. In brief, each point set is converted into a simplicial complex based on a proximity parameter *t*. Recording the appearance and disappearance of topological features over a range of values of *t* produces a persistence diagram. In this work, we used the Euclidean distance as proximity measure between points in a set and compute 0- and 1-dimensional homology groups from Vietoris-Rips simplicial complexes and persistence diagrams for *t* ranging from 0 to half the size of the largest structure using the R package TDA (Fasy et al., 2014). Thus, each 3D SMLM particle for a given protein of interest is represented by a persistence diagram. Finally, to measure topological similarity between nuclear pores, we computed the sliced Wasserstein distance between the corresponding persistence diagrams (Carrière et al., 2017). Clustering was then performed by hierarchical clustering with Ward’s criterion applied to the resulting distance matrix. Clusters were extracted by cutting the resulting dendrogram at a fixed height and each cluster was assigned the label of the most abundant Nup represented in it. To cluster individual Nups for averaging, the subtree induced by each Nup was extracted and cut to generate 5 clusters. Clusters with fewer than 100 members were not processed further. For RANBP2, this produced a single large cluster with more than 100 members which was then further divided into two clusters.

---

**Figure S1.**
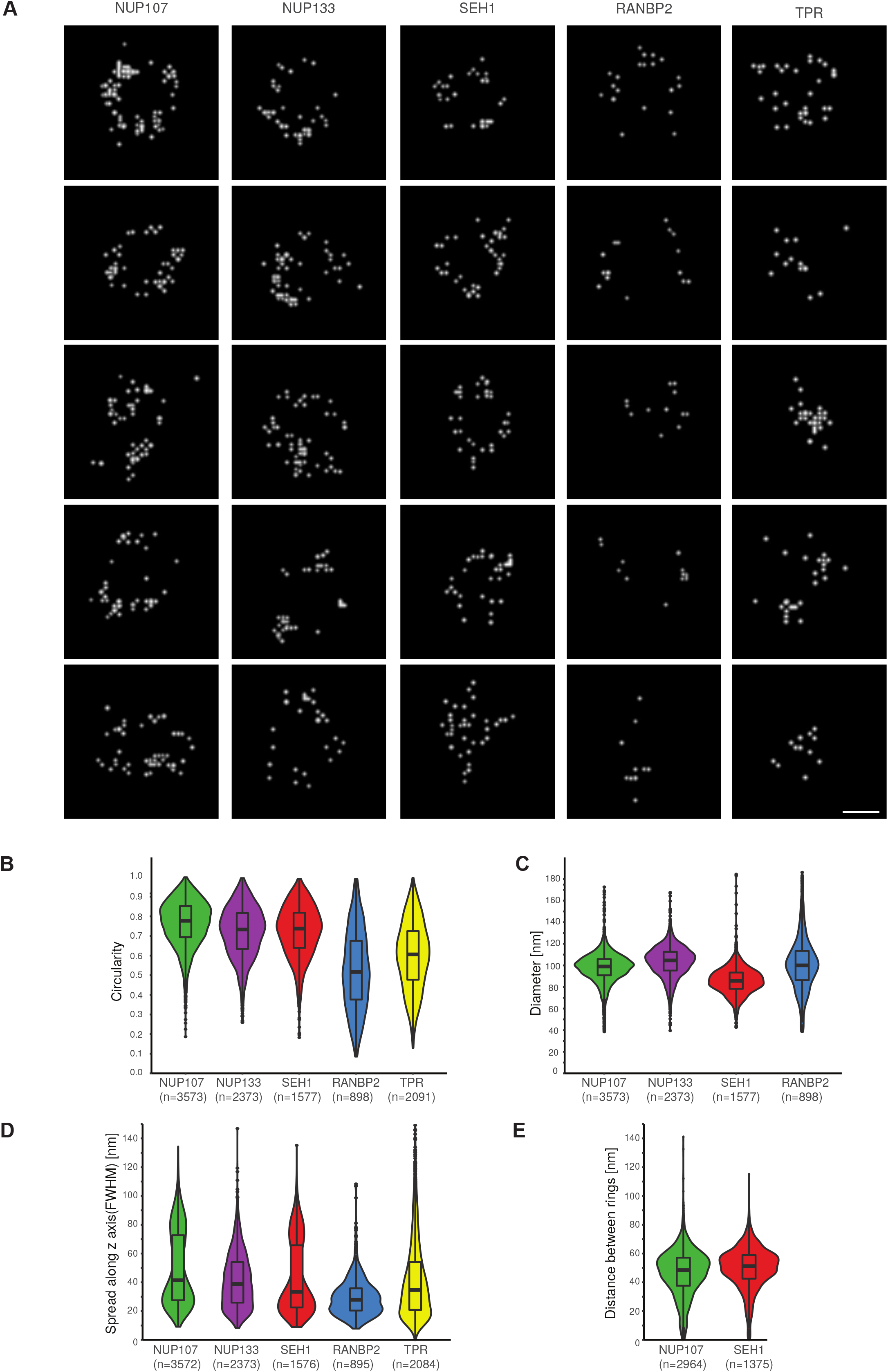
Structural variability within the NPC. **A)** Top view of individual nuclear pores labelled on the different Nups. Scale bar: 5 nm. **B)** Distribution of circularity measurements from individual NPCs for different Nups. **C)** Distribution of diameter measurements from individual NPCs for different NUPs. **D)** Distribution of axial spread measurements from individual NPCs for different Nups. **E)** Distribution of inter-ring distance measured on individual NPCs for NUP107 and SEH1. Number of particles used for this analysis was indicated under each violin plot. RANBP2: light blue, SEH1: red, NUP107: light green; NUP133: purple and TPR: yellow.

**Figure S2.**
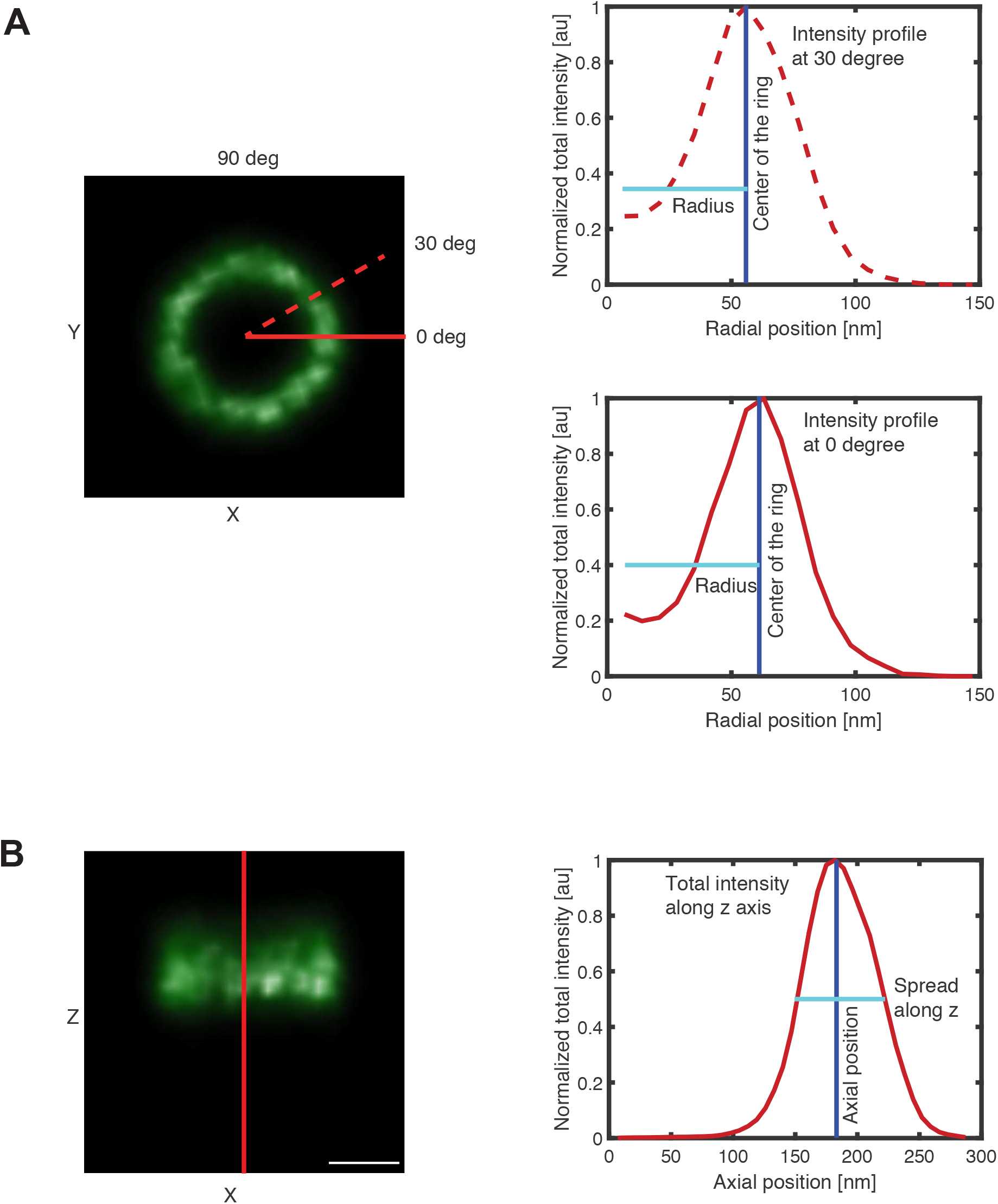
Illustration of radial and axial measurements. **A**) Top view of intensity rendered average density map of the NUP133 (left image). The red lines indicate locations where two radial intensity profiles were calculated (solid line at 0° and dashed line at 30°). Radial intensity profiles (right) along the lines in the image on the left. The blue line indicates the center of the ring obtained through Gaussian fitting. The cyan line indicates the radius. **B)** Side view of the image shown in (A)(left). The red line indicates the direction in z and the corresponding axial intensity profile is shown in the plot (right). The blue line indicates the center of the ring and the spread in z (cyan line) was calculated using full width at half maximum (FWHM). Scale bar, 50 nm.

**Table S1.**
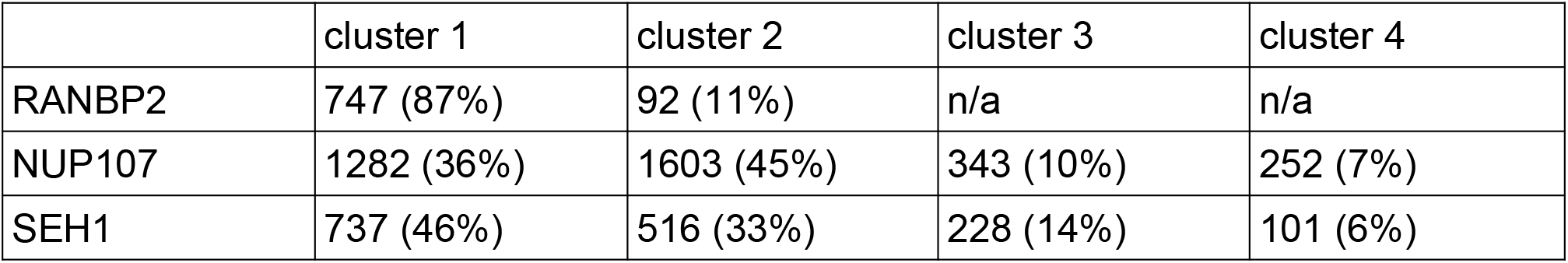
Number of pores in each cluster derived from the induced subtrees. Numbers in parentheses indicate fractions of the total for the corresponding Nup. Totals are 862 pores for RANBP2, 3582 for NUP107 and 1585 for SEH1.

